# L-type calcium channel blocker increases VEGF concentrations in retinal cells and human serum

**DOI:** 10.1101/2021.12.21.473644

**Authors:** Anmol Kumar, Stefan Mutter, Erika B. Parente, Valma Harjutsalo, Raija Lithovius, Sinnakaruppan Mathavan, Carol Forsblom, Markku Lehto, Timo P. Hiltunen, Kimmo K. Kontula, Per-Henrik Groop

## Abstract

**Objective:** Vascular endothelial growth factor (VEGF) plays a key role in diabetic retinopathy (DR). L-type calcium channel blockers (LTCCBs) have been widely used as antihypertensive medication (AHM), but their association with VEGF and DR is still unclear. Therefore, we explored the effect of LTCCBs compared to other AHMs on VEGF concentrations in retinal cells and human serum. Furthermore, we evaluated the association between the use of LTCCBs and the risk of severe diabetic eye disease (SDED).

**Research design and methods:** Müller cells (MIO-M1) were cultured as per recommended protocol and treated with LTCCBs and other AHMs. VEGF secreted from cells were collected at 24 hours intervals. In an interventional study, 39 individuals received LTCCBs or other AHM for four weeks with a four-week wash-out placebo period between treatments. VEGF was measured during the medication and placebo periods. Finally, we evaluated the risk of SDED associated with LTCCB usage in 192 individuals from the FinnDiane Study in an oberservational setting.

**Results:** In the cell cultures, medium VEGF concentration increased time-dependently after amlodipine (p<0.01) treatment, but not after losartan (p>0.01), or lisinopril (p>0.01). Amlodipine, but no other AHM, increased serum VEGF concentration (p<0.05) during the interventional clinical study. The usage of LTCCB was not associated with the risk of SDED in the observational study.

**Conclusions:** LTCCB increases VEGF concentrations in retinal cells and human serum. However, the usage of LTCCBs does not appear to be associated with SDED in adults with type 1 diabetes.

## Introduction

Diabetic retinopathy (DR) is a devastating complication among individuals with diabetes and may lead to permanent vision loss. Given the increasing number of individuals with diabetes worldwide, the prevalence of DR is expected to rise significantly in the coming years (1). Hyperpermeability, hypoperfusion, and neo-angiogenesis of the intraocular microvasculature are key hallmarks of DR and will ultimately lead to anatomical and pathophysiological alterations (2).

Vascular endothelial growth factor (VEGF) is a well-known angiogenic factor in the eye, and is also a major pharmacological target for the treatment of severe and proliferative diabetic retinopathy (PDR) (3, 4). Although, intraocular injections of anti-VEGF antibodies have emerged as a novel and effective pharmacological treatment, long-term efficacy and systemic safety data are lacking, stressing the need to explore alternative pathways and molecular targets for intervention (5). Recently, we reported mutations in the *CACNB2* gene encoding the L-type calcium channel’s β-subunit. These mutations contributed to the severity of PDR. Furthermore, after down-regulating the β-subunit coding *CACNB2* gene by RNA interference, the concentration of VEGF was significantly reduced in retinal cells (6).

Notably, a network meta-analysis showed that renin-angiotensin-aldosterone system inhibitors (RAASi) were associated with reduced risk of progression and increased the possibility of regression of DR (7). However, although the calcium channel blockers seemed to worsen the outcome compared to placebo, it was not statistically significant (7). L-type calcium channel blockers (LTCCB) have been widely prescribed to treat arterial hypertension as monotherapy or in combination with other classes of antihypertensive medications (AHM) such as angiotensin-converting enzyme inhibitors (ACEi), angiotensin II receptor blockers (ARB), β blockers and diuretics (8, 9). However, the relationship between LTCCB and DR is still poorly understood.

Therefore, in this work, we aimed to explore the effect of LTCCBs in comparison to other AHMs on the VEGF concentrations in retinal cell culture and serum of human subjects. We used human retina-derived Müller cells for *in vitro* experiments as these cells are a crucial source of VEGF in the pathogenesis of DR (10, 11). Additionally, we evaluated the effects of different classes of AHMs on the serum VEGF concentrations in a clinical trial setting (The GENRES Study). Finally, we evaluated the association between the use of LTCCB and the risk of severe diabetic eye disease (SDED) in adults with type 1 diabetes from the Finnish Diabetic Nephropathy Study (FinnDiane) cohort.

## Research Design and Methods

### Retinal cell culture study

Müller glia or Müller cell lines, MIO-M1 were obtained from Prof. Astrid Limb (12). Cells were grown in Dulbecco’s Modified Eagle Medium (41965-039, Gibco/life technology), 10% Fetal Bovine Serum (10270106, Gibco/life technology), Penicillin-Streptomycin (15140122, Gibco/life technology) and Normocin™ (ant-nr-1, InvivoGen) in a humidified 5% CO_2_ cell culture incubator at +37°C temperature. Cells were plated in 6-well plates (140675, Nunc™) with seeding density (~10^6^ cells/well). Three medications - amlodipine besylate (A5605, Sigma-Aldrich), losartan potassium (phr1602-1g, Sigma-Aldrich) and lisinopril (phr1143-1g, Sigma-Aldrich) were dissolved in solvents as per the manufacturer’s recommendation. The medications and the dimethyl sulfoxide (DMSO)/1X Phosphate-buffered saline (PBS) control (the solvent used to dissolve the medication) were added in equal volume to the cell culture medium. The cell culture medium containing medications and control in equal volume were changed every 24 hours until the end of the experiment. Cell culture medium was collected and centrifuged at +8°C temperature at 3000 rpm for 5 minutes to remove cell debris, and the resulting supernatants were collected and stored in a −80°C freezer for further analysis.

### Interventional clinical study

The GENRES Study is a randomized, double-blind, placebo-controlled, crossover study, which included Finnish men with moderate hypertension but without drug-treated diabetes. The detailed protocol can be found in a previous publication (13). In brief, the GENRES study started with a four-week run-in placebo period, before which the participants discontinued their previous antihypertensive medication, in case they were on medication. Then, participants were randomized into different sequences of four-week treatment periods taking one of the AHM (amlodipine, bisoprolol, hydrochlorothiazide, losartan) at a time. Thus, each participant underwent four separate monotherapy periods, separated by four-week placebo periods (13). Serum VEGF concentration was measured at seven different time points, at the end of the three placebo and the four drug treatment periods. The serum at the end of the first placebo period was not included in the analysis, since the blood sample was taken after an overnight fast as opposed to the other blood samples, which were taken in the non-fasting state.

### VEGF measurements

VEGF measurements were performed double-blinded in serum and cell culture media using the Human VEGF Quantikine ELISA Kit (DVE00 & SVE00, R&D systems/biotechne) according to the manufacturer’s protocol.

### Observational cohort study

The outcome was SDED, defined as a composite outcome including proliferative diabetic retinopathy (PDR), initiation of laser treatment or anti-VEGF therapy, diabetic maculopathy, vitreous hemorrhage and vitrectomy identified from the Finnish Care Register for Health Care until the end of 2015.

Information on purchases of AHMs was obtained from the Finnish Drug Prescription Register (maintained by the National Social Insurance Institution since 1994, which contains information on all prescribed, purchased, and reimbursed medications in outpatient care) until the end of 2015. Medications were coded according to the Anatomic Therapeutic Chemical (ATC) classification system. The renin-angiotensin-aldosterone system inhibitors (RAASi) were identified by an ATC code starting with C09 including both ACEis and ARBs. To identify individuals on RAASi monotherapy, we excluded individuals with a prescription record of combination products that contained other AHMs in the same pill (coded with C09B and C09D) or with a prescription for any other AHMs prior to the baseline or during the follow-up. LTCCBs in the register included the following ATC codes: C07FB02, C07FB03, C08CA01, C08CA02, C08CA03, C08CA05, C08CA07, C08CA10, C08CA13, C09BB02, C09BB04, and C09BB05. Refill adherences were calculated using the so-called proportion of days covered method with a prescription refill interval of 90 days.

For this longitudinal study, 192 individuals were selected from the FinnDiane study, which is an ongoing, nationwide, multicenter cohort of adults with type 1 diabetes in Finland (14, 15). Type 1 diabetes was defined as age of diabetes onset below 40 years and requiring insulin within the first year after diagnosis. Currently, more than 5,400 individuals have undergone a detailed clinical examination at their first FinnDiane visit. We excluded 1,933 individuals due to SDED at baseline. These data (history of laser treatment) were registered based on medical records at baseline visit and obtained by the attending physician or nurse using a standardized form. The selection criteria for the remaining 3,468 individuals can be found in the Supplementary Figure 1. First, we required the study visit to take place at least half a year before the end of the prescription records at the end of 2015 and no SDED in the first half year of follow-up. As shown in Supplementary Figure 1, our cohort is made up of two groups: those on LTCCB plus RAASi and other AHM and a control group of those on RAASi as monotherapy. In both groups individuals had been on RAASi therapy at baseline, defined as either already being on RAASi prior to the study visit or starting RAASi therapy no longer than half a year after the visit. We further required at least 2 purchases of RAASi during the follow-up. The same selection criteria were applied for LTCCB in the group LTCCB plus RAASi and other AHM. In this latter group, individuals were allowed to use other AHMs, because, according to the hypertension guidelines, LTCCBs are usually considered as the second or third line drug after RAASi (16, 17). Due to the strict adherence to the guidelines in Finland, we did not find enough individuals using LTCCB as monotherapy (n = 11) to be able to investigate them as a separate group.

### Statistical analysis

Cell culture data were analyzed using unequal variance student’s t-test. A p-value below and equal to 0.01 was considered significant.

In the clinical study (The GENRES study), the mean VEGF concentration of the three placebo periods was used as the baseline for each subject. Since VEGF concentrations at baseline and changes after the drug treatment periods were non-normally distributed, the nonparametric Wilcoxon signed ranks test was used to test the statistical significance of the drug-induced changes. A p-value below 0.05 was considered significant.

In the observational study, a multivariable logistic regression analysis was used to study the associations between the usage of LTCCB plus RAASi and AHM (n=62) and the incidence of SDED as the outcome. Since RAASi usage has shown an association with reduced risk of progression and increased possibility of regression of DR (7), first, we performed a logistic regression using the adherence to RAASi usage as the independent variable and the incidence of SDED (n=130) as the outcome. Since the adherence to RAASi usage was associated with the outcome, this variable was included as a covariate in the logistic regression model for the analysis of LTCCB plus RAASi and AHM and incident SDED. Clinical baseline variables significantly different at baseline between those that developed SDED and those that did not, i.e., age, age of diabetes onset, albuminuria and HbA1c were used as covariates alongside the adherence to RAASi treatment during follow-up, other antihypertensive medication on top of LTCCB plus RAASi during follow-up, and the presence of any retinopathy other than SDED at baseline. For continuous variables, p-values were calculated using t-tests for normally distributed variables and Mann-Whitney U-tests for non-normally distributed variables. For categorical variables, we used the χ^2^-test. A p-value below 0.05 was considered significant. All analyses on observational data were performed in R version 3.6 (18).

## Results

### Retinal cell culture study

After treating the MIO-M1 cells with 10 μM amlodipine, the secreted VEGF protein concentration increased in the medium at all time points of the treatment (24, 48, 72 and 96 hours), compared to the control DMSO/1X PBS (Figure 1). In contrast, VEGF did not change after lisinopril or losartan treatment, except a slight increase at 96 hours of the losartan treatment (Figure 1). Additionally, we tested lower concentrations of amlodipine (1 μM and 5 μM), but they did not affect the VEGF concentration in the medium of the cell culture at 24, 48, 72 and 96 hours (Supplementary Figure 2). Finally, we performed cell viability tests at the end of our experiment (96 hours). Amlodipine (10 μM) reduced cell proliferation, whereas losartan (10 μM) and lisinopril (10 μM) did not affect the cell viability or proliferation compared to the control (DMSO/1XPBS) (Supplementary Figure 3).

**Figure 1:**
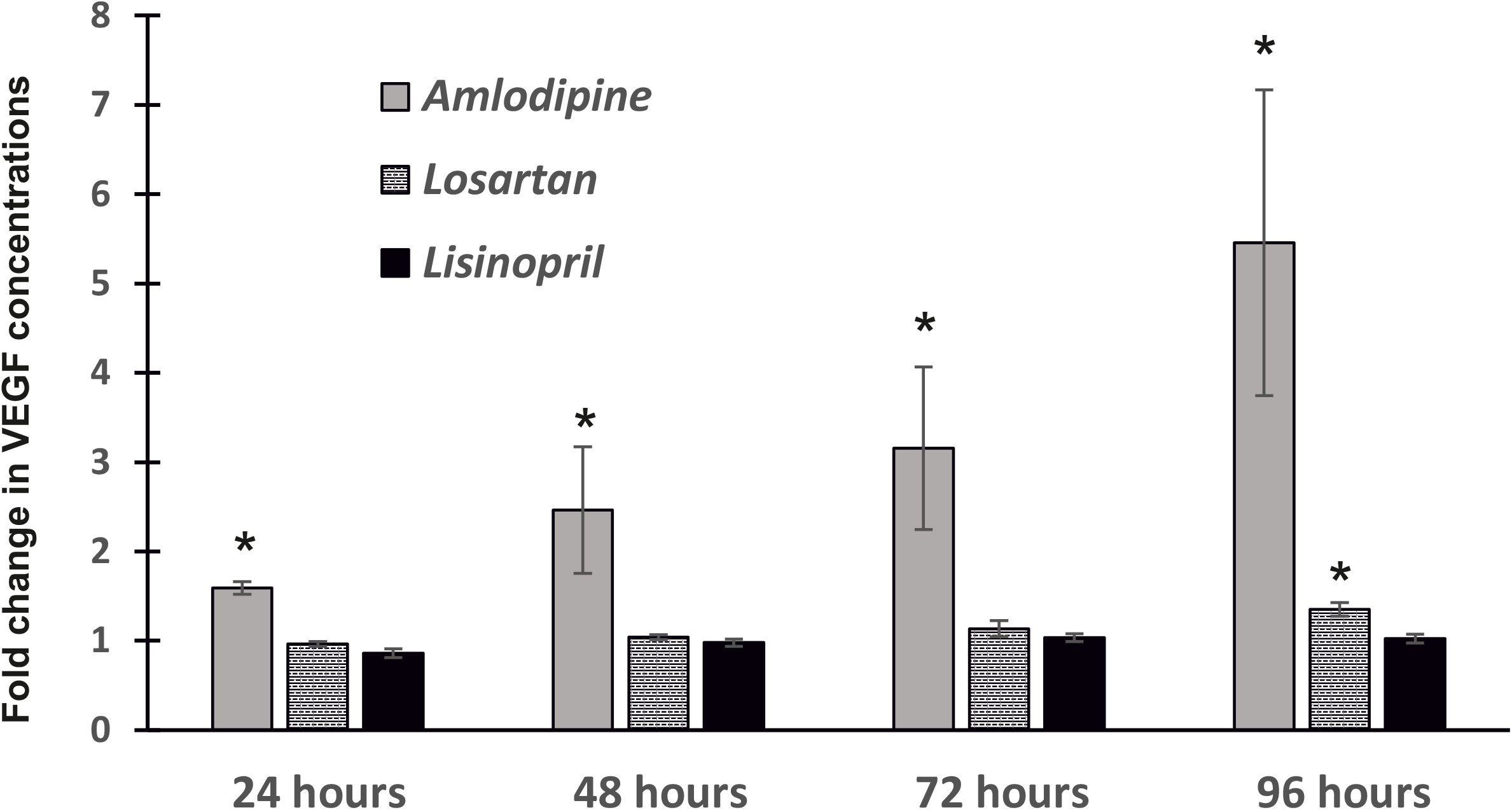
Mean fold changes in the VEGF concentrations in MIO-M1 cell culture medium after treatment with amlodipine (10 μM), losartan (10 μM) and lisinopril (10 μM) at different time points (24 hours, 48 hours, 72 hours and 96 hours) calculated from two independent experiments each with triplicates for treatments. The fold changes in VEGF concentrations are calculated to control treatment with solvent (DMSO and 1X PBS) of each drug at each time point. All statistical analyses have been done between drug treatments and respective control (DMSO/1XPBS) treatment at each time point. *p value < 0.01.

### Interventional clinical study

During the placebo periods, the median serum VEGF concentration was 277 pg/mL in 39 men with hypertension. During the LTCCB treatment period, there was a median increase of 18 pg/ml in the serum VEGF concentration (p=0.03) (Figure 2). Treatment with hydrochlorothiazide induced a nonsignificant increase (p=0.09) while treatment with losartan a nonsignificant decrease (p=0.25) in the VEGF concentration (Fig. 2).

**Figure 2:**
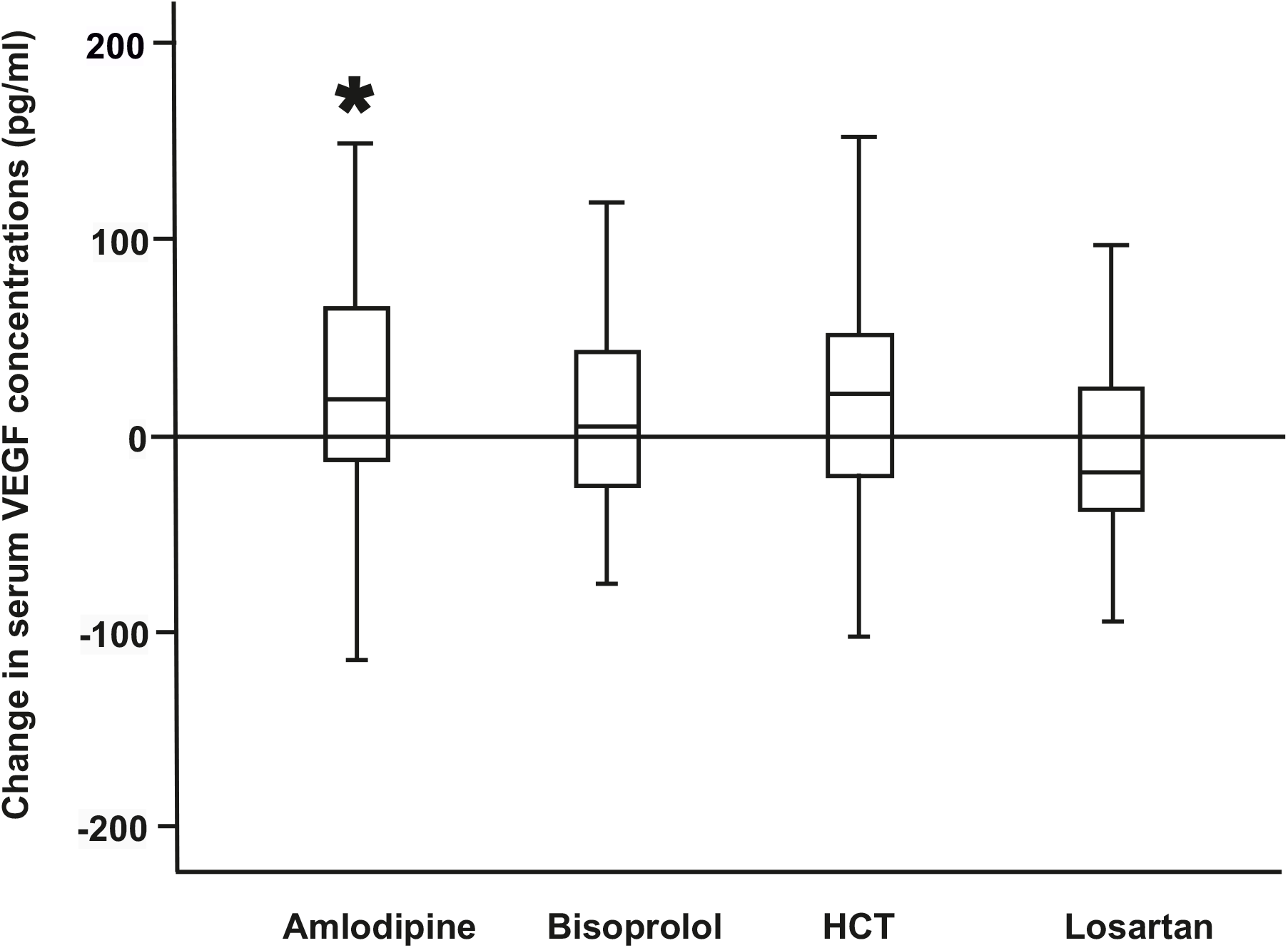
Changes in serum VEGF levels after 4 weeks of daily monotherapies with amlodipine (5 mg), bisoprolol (5 mg), hydrochlorothiazide (25 mg) and losartan (50mg). N=39. *p value < 0.05.

### Observational study

During a mean follow up of 8.6 ± 5.7 years, 70 events of SDED occurred. Individuals who developed SDED had a younger age at onset of diabetes and higher glycated hemoglobin (HbA1c) at baseline than those that did not develop SDED. The clinical characteristics of the individuals at baseline, according to the incidence of SDED, are shown in Table 1. The higher refill adherence to RAASi was associated with a lower risk of SDED (OR 0.97, 95%CI [0.95-0.99], p=0.005) in an unadjusted model. The use of LTCCB plus RAASi and other AHM was neither associated with increased odds of SDED (OR 0.94, 95% CI [0.50-1.77], p=0.85) in the unadjusted model nor in the adjusted model (OR 6.12, 95% CI [0.88-42.45], p=0.07).

**Table 1.** Characteristics of individuals with type 1 diabetes participating in the observational study.

|  | **No SDED** | **Incident SDED** | **P** |
| --- | --- | --- | --- |
| N | 122 | 70 |  |
| LTCCB (%) | 32.8 | 31.4 | 0.97 |
| Other AHM (%) | 31.1 | 22.9 | 0.29 |
| Women (%) | 41.0 | 38.6 | 0.86 |
| Age (years) | 42.82 ± 10.65 | 36.30 ± 11.93 | <0.001 |
| Age of diabetes onset (years) | 18.34 ± 8.10 | 12.92 ± 7.72 | <0.001 |
| Diabetes duration (years) | 24.50 ± 11.75 | 23.38 ± 9.71 | 0.50 |
| Systolic blood pressure (mmHg) | 141 ± 18 | 137 ± 17 | 0.10 |
| Diastolic blood pressure (mmHg) | 80 ± 10 | 81 ± 10 | 0.43 |
| Total cholesterol (mmol/L) | 4.00 ± 1.12 | 4.89 ± 1.04 | 0.93 |
| Triglycerides (mmol/L) | 0.03 (0.69, 1.33) | 1.08 (0.81, 1.40) | 0.08 |
| eGFR (mL/min/1.73m<sup>2</sup>) | 93 ± 28 | 96 ± 29 | 0.43 |
| Albuminuria (%) | 31.1 | 70.0 | <0.001 |
| HbA<sub>1c</sub> (%) | 8.2 ± 1.3 | 9.3 ± 2.0 | <0.001 |
| HbA<sub>1c</sub> (mmol/mol) | 66 ± 14 | 78 ± 22 | <0.001 |
| RAASi before baseline (years) | 3.04 ± 2.77 | 2.79 ± 2.31 | 0.53 |
| Adherence to RAASi (%) | 74.94 ± 17.20 | 70.05 ± 20.58 | 0.08 |
| Any retinopathy except SDED (%) | 52.6 | 67.7 | 0.07 |

Data are mean ± SD or median (interquartile range). Other AHM: antihypertensive medication other than LTCCB or RAASi.

## Discussion

In the present study, we showed that amlodipine, an LTCCB, increases the secretion of VEGF *in vitro* by cells of human retinal origin, in a time-dependent manner. In addition, we showed that four weeks of amlodipine usage also increases the serum VEGF concentration in individuals with arterial hypertension. However, in our observational study, the use of LTCCB was not associated with an increased risk of SDED in adults with type 1 diabetes.

Our observations are supported by a previous report showing that nifedipine, a first-generation LTCCB, induces VEGF secretion in human coronary artery smooth muscle cells (HCSMCs) [16]. We used 24-96 hours of treatment in contrast to 18-24 hours in previous *in vitro* studies [16, 17]. This prolonged *in vitro* treatment may better represent a chronic pharmacological intervention (19). Additionally, we observed an anti-proliferative effect of amlodipine on MIO-M1 cells, similar to what has been shown with nifedipine on HCSMCs (19). Another study has reported no effect of amlodipine on VEGF secretion by HCSMCs. However, the shorter treatment duration and lower medication dosage (5 μM) used in that study may explain the observed differences compared to our results (20). To reinforce this hypothesis, we also tested 1 μM and 5 μM of amlodipine on MIO-M1 cells, and we did not find any changes in the VEGF concentration at any time of the treatment (24-96 hours). Importantly, DR disrupts the blood-retinal barrier increasing perfusion of medications into the ocular compartment, thus altering intraocular pharmacokinetics and diffusion of medication molecules (21, 22). This may increase the diffusion of L-type calcium channel blockers from the systemic circulation into the ocular compartments in individuals with DR and thereby elevating their intraocular concentrations. Our data from the interventional clinical study show that the serum VEGF concentration increases after four weeks of amlodipine treatment (5 mg/day). Although a previous study reported a positive association between serum and vitreous VEGF levels (23, 24), we are not able to conclude from our results whether an elevated serum VEGF concentration also translates into an increased VEGF concentration in the vitreous humour. Furthermore, it is not possible to ensure that the increased serum VEGF concentration after four weeks of amlodipine usage will persist after years of treatment. Further *in vivo* experiments using animal models are needed to clarify these questions, since collecting vitreous samples from human subjects is challenging.

The observational study showed that long-term usage of LTCCB plus RAASi and other AHM was not associated with SDED. We are not sure whether the lack of association is due to the combination of LTCCB with RAASi, since RAASi have been linked to lower risk of DR (7, 9). However, we included refill adherence to RAASi therapy as a covariate in our models. In our study, it was not possible to evaluate the risk of SDED related to the use of LTCCB as monotherapy, because the number of individuals using LTCCB without other medication in the FinnDiane cohort was too small. Indeed, this is aligned with the hypertension treatment guidelines in people with diabetes (16, 17) in which the calcium channel blockers are not the first-line choice and are commonly prescribed in association with RAASi as second or third-line therapy. Although we did not find an association between the use of LTCCB and SDED, we found that a higher adherence to RAASi was associated with lower odds of SDED, similar to what has been previously shown (7).

The major strength of the current study is the *in vitro* experiments with retina-derived Müller cells showing for the first time a time-dependent effect of LTCCB on VEGF concentrations. Additionally, we validated these *in vitro* findings using a unique interventional clinical trial that showed an increase in serum VEGF after four weeks of exposure to LTCCB. The main drawback of our study is that we were not able to measure VEGF in the vitreous humour to evaluate the correlation between the serum and the vitreous VEGF concentrations. Furthermore, despite the inclusion of one of the largest cohorts of individuals with type 1 diabetes (the FinnDiane Study), there was not a large enough number of individuals using LTCCB as monotherapy in order to be able to explore a potential adverse association between the use of LTCCB and SDED.

## Conclusion

Our results show that LTCCB increases VEGF concentrations in retinal origin cells and in human serum, but its usage in combination with RAASi and other AHM does not seem to be associated with SDED in adults with type 1 diabetes in an observational setting. Further studies including a larger sample of individuals with other types of diabetes using LTCCB as monotherapy, as well as studies with animal models to measure the VEGF in the vitreous humour are warranted to evaluate the impact of chronic use of LTCCBs on the development and progression of DR.

## Acknowledgements

The skilled technical assistance by Heli Krigsman and Susanna Saarinen, is gratefully acknowledged. The authors also acknowledge the participants, physicians and nurses at FinnDiane study centers (see supplemental table 1). Parts of this study have been presented in abstract form at the 56^th^ Annual Meeting of the European Association for the Study of Diabetes.

## Author’s contribution

AK conceived the project, executed experiments, analyzed data, wrote the manuscript. SM and EBP equally contributed to processing, analysing the observational data and writing the manuscript. TH analysed interventional data. VH, RL, SiMa and CF contributed to collection and processing of data. ML, KK and PHG supervised the studies. All authors interpreted the results as well as reviewed and approved the final version of the manuscript. PHG is the guarantor of *in vitro* and observational study work, and, as such, had full access to all the data in those studies and takes responsibility for the integrity of those data and the accuracy of those data analyses. KK is the guarantor of the interventional study work and, as such, had full access to all the data in this study and takes responsibility for the integrity of those data and the accuracy of this data analysis.

---

**Supplementary Figure 1:**
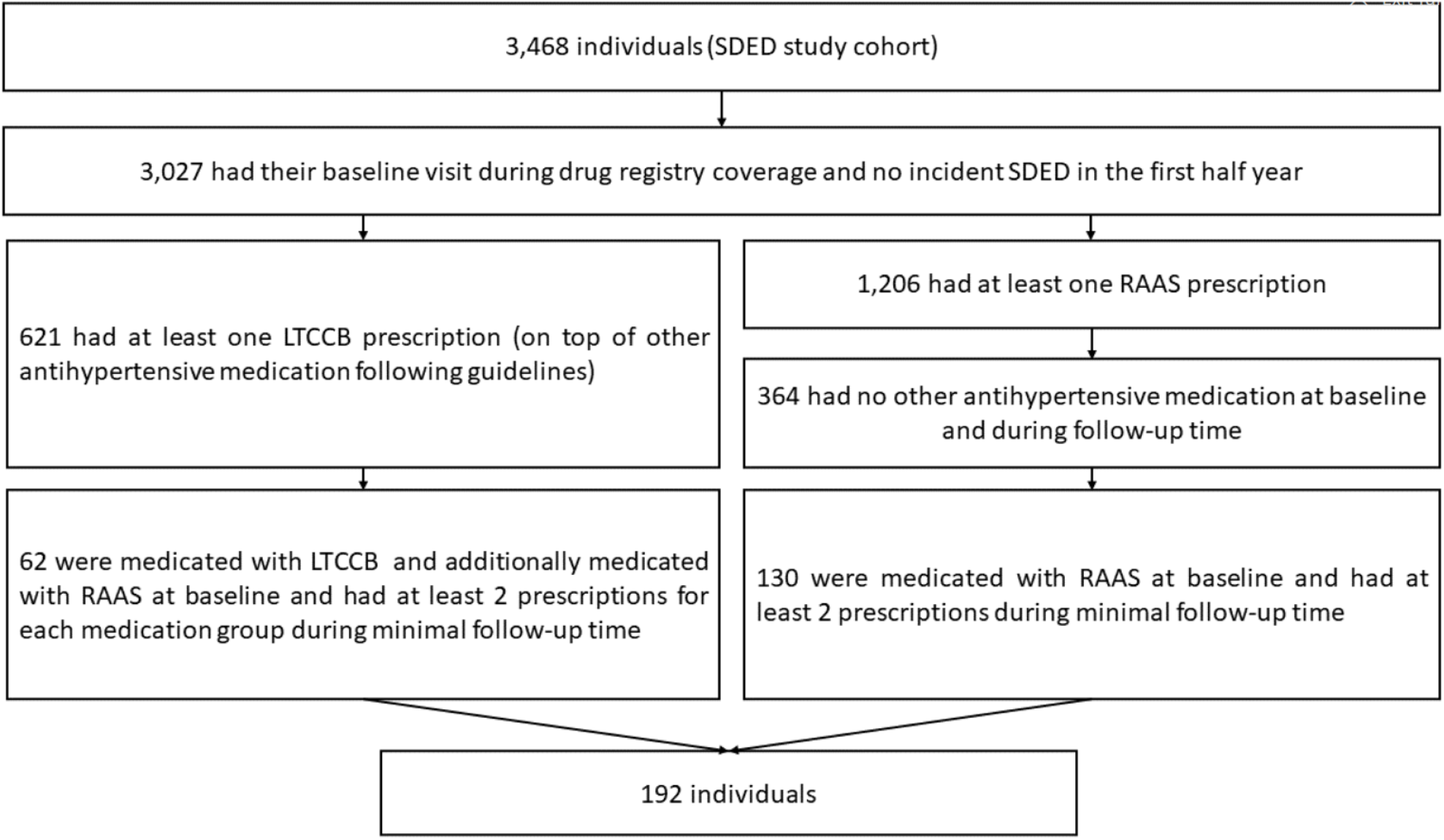
Schematic diagram depicting selection criteria in the observational study with FinnDiane cohort.

**Supplementary Figure 2:**
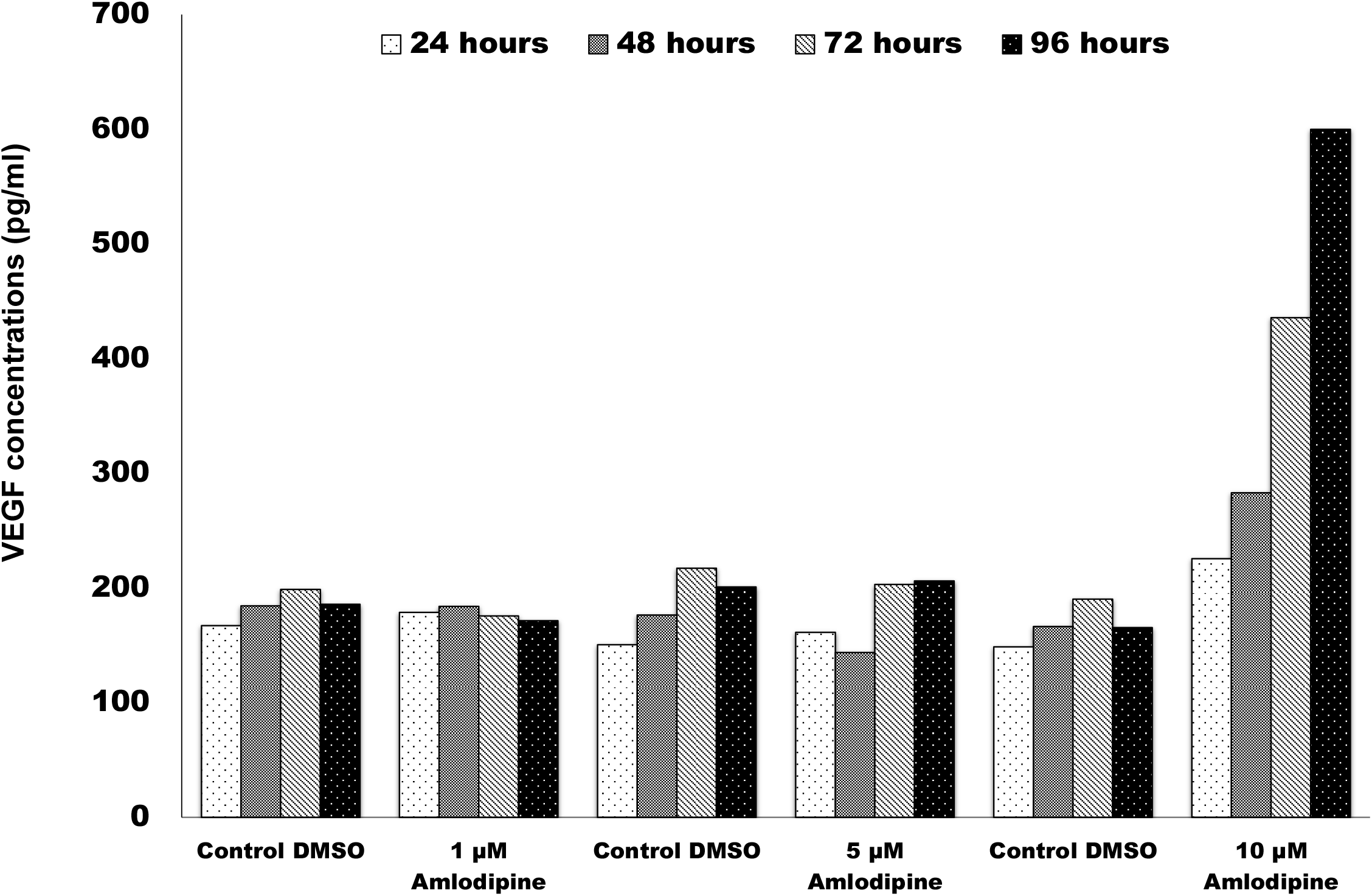
Dose response (1, 5 and 10 μM) experiment for determination of effects of amlodipine on VEGF secretion in MIO-M1 cells at different concentrations and time points. Values for VEGF concentrations are calculated as average of 3 wells for every treatment from a single experiment.

**Supplementary Figure 3:**
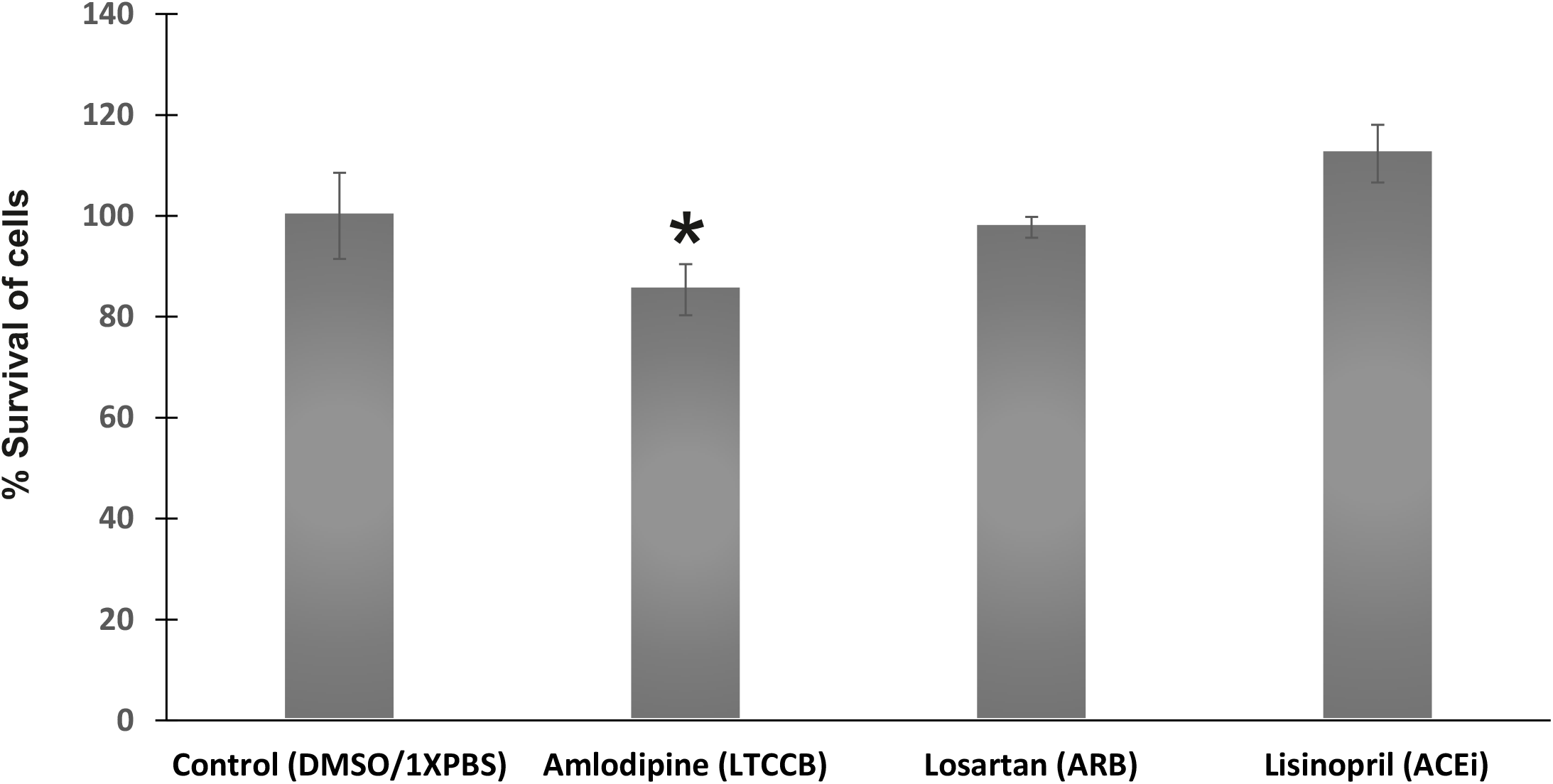
Survival of MIO-M1 cells at end of drug treatment (96 hours). Survival mean percentages are calculated from two independent experiments.*p value = 0.01.

